# Aging associated altered response to intracellular bacterial infections and its implication on the host

**DOI:** 10.1101/2020.09.15.298158

**Authors:** Sheryl E. Fernandes, Alakesh Singh, R.S. Rajmani, Siddharth Jhunjhunwala, Deepak K. Saini

## Abstract

The effects of senescence and aging on geriatric diseases has been well explored but how these influence infections in the elderly have been scarcely addressed. Here, we show that several innate immune responses are elevated in senescent cells and old mice, allowing them to promptly respond to bacterial infections. We have identified higher levels of iNOS as a crucial host response and show that p38 MAPK in senescent cells acts as a negative regulator of iNOS transcription. In old mice, however the ability to impede bacterial proliferation does not correlate with increased survival as elevated immune responses persist unabated eventually affecting the host. The use of anti-inflammatory drugs that could consequently be recommended also decreases iNOS disarming the host of a critical innate immune response. Overall, our study highlights that infection associated mortality in the elderly is not merely an outcome of pathogen load but is also influenced by the host’s ability to resolve inflammation induced damage.

**Summary statement:** Using cellular models and old mice we demonstrate the effect of aging on host response to bacterial infections. Aged systems mount a more effective anti-bacterial innate immune response but its persistence results in mortality of the host.

## Introduction

Organismal aging is defined as a gradual loss in physiological function and homeostasis (Flatt, 2012) which predisposes the elderly to several diseases. In 1881, the evolutionary biologist A. Weissman proposed that the functional decline associated with aging was likely to be a result of the finite capacity of cells in the body to divide (Childs *et al*., 2017). This remained a theory till much later when Leonard Hayflick for the first time demonstrated that indeed, cells from the body can divide only a finite number of times before reaching a non-dividing state (Hayflick and Moorhead, 1961; Hayflick, 1965), which is now referred to as cellular senescence. Several studies since then have shown that senescent cells accumulate in several tissues with age (Dimri *et al*., 1995; Jeyapalan *et al*., 2007) and their targeted removal not only increases lifespan but also ameliorates common aging associated pathologies (Baker *et al*., 2011; Campisi and Deursen, 2016; Jeon *et al*., 2017).

Numerous mechanisms are known to trigger senescence of cells in the body like telomere attrition, DNA damage and oxidative stress due to mitochondrial dysfunction (Ben-Porath and Weinberg, 2005; Hernandez-segura, Nehme and Demaria, 2018). However, despite the multiple triggers, the eventual outcome is a persistent DNA damage response (DDR) that activates cell cycle inhibitors and initiates senescence (Rossiello *et al*., 2014; Salama *et al*., 2014). Therefore, genotoxic agents (e.g. Ionizing radiation, BrdU and Doxorubicin) are popularly used to induce and study senescence *in vitro* (Chen, Ozanne and Hales, 2007; Petrova *et al*., 2016; Wang, Boerma and Zhou, 2016).

After committing to this state of permanent cell cycle arrest, many cellular changes occur altering the signalling landscape resulting in cells which are phenotypically distinct from their non-senescent counterparts (Salama *et al*., 2014). The secretome of senescent cells, referred to as senescence associated secretory phenotype (SASP) consists of a variety of growth factors, cytokines and chemokines. SASP has been linked to a plethora of geriatric metabolic and degenerative disorders (Baker *et al*., 2011; Childs *et al*., 2017), however, senescence and its implication on infection has not yet been well studied.

Old individuals do not respond similarly to young infected ones clearly indicating that an aged system perceives pathogens differently. Instead of expected clinical symptoms like fever, atypical symptoms like nausea or weight loss are observed making diagnosis challenging as the symptoms are often attributed to other age-related co-morbidities (Byng-Maddick and Noursadeghi, 2016). Therefore, understanding the interplay between aging and infection at cellular and organismal level becomes important.

In this study, we induce senescence in an established epithelial cell model of infection to understand the impact of aging on infection of a well-studied intracellular pathogen, *Salmonella* Typhimurium. Epithelial cells are important targets for intracellular pathogens and play a crucial role in deciding pathogenesis and prognosis of disease. However, their critical immune function in pathogen defense is only beginning to be understood (Krausgruber *et al*., 2020).

Here, we show that senescent epithelial cells are better at inhibiting intracellular bacterial proliferation and investigation of anti-pathogen cellular mechanisms reveal that innate immune responses in senescent cells are significantly up regulated compared to non-senescent cells. With the help of molecular inhibitors, we identify elevated levels of Nitric oxide (NO) as the most important modulator of infection and show that p38 MAPK in senescent cells acts as a negative regulator of NO. Later, *in-vivo* studies of infection in naturally aged BALB/c mice shed light on how simultaneous senescence in multiple organ systems affect pathogen dissemination and infection. We also report lesser bacterial burden in old mice compared to young mice but interestingly no significant differences in lethality. This we demonstrate is an outcome of old mice limiting bacterial infection more competently due to a higher nitrosative response but an inability to upregulate host protective responses to infection induced inflammation. Consequently, the use of anti-inflammatory drugs maybe advocated, which also helps to reduce disease severity of co-morbidities driven by SASP (Crofford, 2013). However, we show that several clinically approved and over-the-counter (OTC) anti-inflammatory drugs may also perturb innate immune responses of senescent cells therefore restricting their usefulness during infection.

Our findings are finally validated in an another model of *Mycobacterium tuberculosis* infection of senescent lung epithelial cells and aged mice to overcome the concerns of a cell line or pathogen specific observation and eventually reveal a novel advantage and alternate function of accumulated senescent cells in the elderly.

## Results

### Bacterial proliferation in senescent cells is reduced compared to non-senescent cells

HeLa is widely accepted as an Epithelial cell model system for *Salmonella* infection and many fundamental virulence associated studies have been done using these cells (Hannemann, Gao and Gala, 2013; Bowden *et al*., 2014; Alvarez *et al*., 2017; Aguilar *et al*., 2020). Therefore, we adopted HeLa as the cellular model for our study. We used BrdU, a genotoxic agent, to induce DNA damage and to mimic persistent DDR triggered cellular senescence *in vitro*. BrdU mediated senescence has been previously used to explore senescence associated changes by various groups including for HeLa cells (Masterson and Dea, 2007; Lim *et al*., 2010; Nair, Bagheri and Saini, 2014). Importantly, BrdU is incorporated specifically into the DNA thereby avoiding uncharacterized non-specific effects of other genotoxic agents like Doxorubicin or ionizing radiation (IR) which can also generate free radicals that can oxidize and perturb protein and lipid homeostasis affecting infection directly (Thorn *et al*., 2011; Reisz *et al*., 2014). All these factors made BrdU induced senescence an ideal method to study downstream effects of senescence on infection.

Senescence in BrdU treated HeLa cells was confirmed by Senescence-Associated β-galactosidase (SA β-gal) staining (Fig 1A), a well-accepted physiological marker (Dimri *et al*., 1995). Other senescence associated molecular markers like increase in the expression of p21/ *CDKN1A*, a cell cycle inhibitor and phosphorylation of Chk2, a signalling protein in the DNA damage detection cascade (Chen, Hales and Ozanne, 2007) were also confirmed in the treated cells (Supplementary Fig. S1A and S1B).

**Figure 1.**
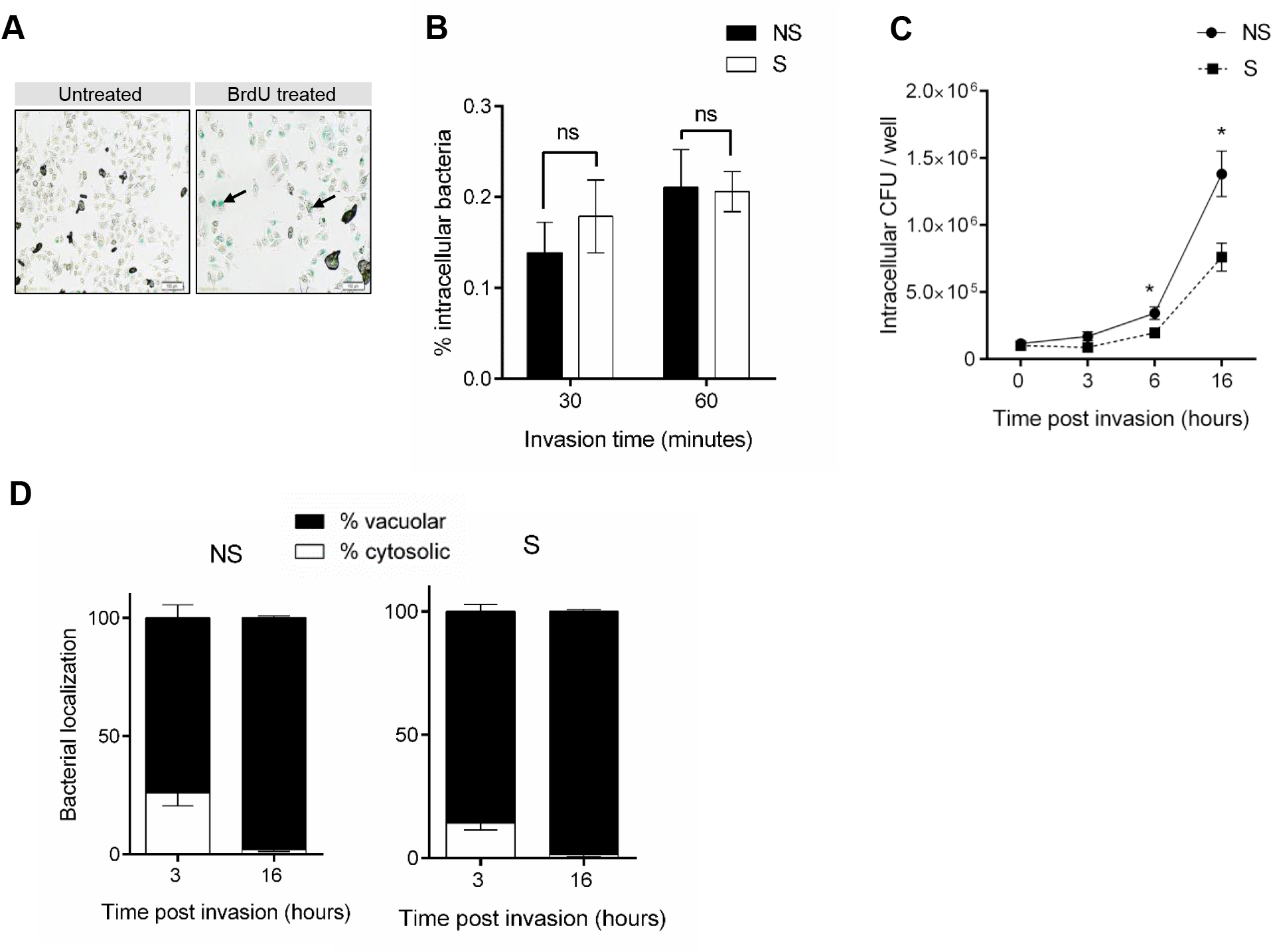
Effect of senescence on *Salmonella* infection. **A**. SA β-gal staining of BrdU treated HeLa cells to confirm senescence. Scale bars, 100 µm; arrows indicate cells positive for staining. **B**. Comparison of *S*.Typhimurium invasion between non-senescent (NS) and senescent (S) cells. Percent intracellular bacteria calculated as (CFU at 30 or 60 min/input CFU) ×100. **C**. Comparison of intracellular bacterial proliferation between NS and S cells. Equal number of host cells were infected at MOI 10. After 1 h, extracellular bacteria were killed by treating with 100 µg/ml Gentamicin for 30 min. Infected cells were then incubated with medium containing 10 µg/ml Gentamicin till the time points mentioned. Bacterial CFU was determined from cell lysates. **D**. Chloroquine assay to determine vacuolar or cytosolic localization of bacteria. Prior to lysis for CFU determination, cells were incubated with 400 µM Chloroquine for 1h (cytosolic CFU) or without Chloroquine (total CFU). The data represents mean ± SEM from at least three independent experiments. Statistical significance of differences was analysed by Mann-Whitney U test, \**P* ≤ 0.05. For all experiments NS, non-senescent and S, senescent.

Once the cellular senescence model was validated, to investigate if bacterial invasion and survival varies between non-senescent (NS) and senescent (S) cells, equal numbers of NS and S HeLa cells were infected with *Salmonella* Typhimurium NCTC 12023 for 30 minutes and 60 minutes. There was no difference in bacterial invasion determined through colony forming units (CFU) on Salmonella Shigella (SS) agar (Fig 1B). Further, to compare bacterial proliferation and survival in NS and S cells, *Salmonella* were allowed to invade host cells for 60 minutes (invasion), after which extracellular bacteria were removed by Gentamicin treatment. Viable intracellular bacteria were then enumerated in terms of CFU immediately at invasion (indicated as 0h post invasion in Fig 1C) and at 3h, 6h and 16h post invasion. At 3h, there was no significant difference in CFU, however, at 6h and 16h, S cells harboured significantly lesser number of bacteria compared to NS cells (Fig 1C), indicating that while cellular senescence does not affect invasion, it impairs bacterial survival and proliferation. The CFU data was corroborated with fluorescence imaging of GFP-HeLa cells infected with *Salmonella* constitutively expressing mCherry (Fig S2). At 16h post invasion, lesser bacterial numbers could be visualized in S cells compared to NS cells. Additionally, a similar reduction in *Salmonella* proliferation was observed in BrdU induced senescent HepG2 hepatocyte cells compared to non-senescent HepG2 cells (Fig S3) indicating that the effect of senescence on *Salmonella* proliferation was consistent across cell lines.

Previously it has been shown that intracellular localization of *Salmonella* can influence bacterial proliferation. Loss in integrity of *Salmonella* containing vacuoles (SCVs) results in their entry into the cytoplasm; and these bacteria proliferate faster than those contained within SCVs (Brumell *et al*., 2002). Having observed lesser bacteria in S cells, we investigated if bacteria were able to escape into the cytosol of NS cells but were restricted to vacuoles in S cells. For this, we determined percent vacuolar and cytosolic bacteria using a well-established Chloroquine assay (Knodler, Nair and Steele-mortimer, 2014). No significant difference in percent cytosolic bacteria was observed at both early (3h) and late (16h) time points of infection and bacteria were found to be majorly vacuolar in both NS and S cells (Fig 1D). This suggested that the difference in CFU is not due to their intracellular localization but due to some other factor intrinsic to senescent cells.

### Many antimicrobial factors are altered in senescent cells

Since S cells showed significantly lower bacterial proliferation (Fig 1C), we hypothesized that antimicrobial factors maybe up regulated in these cells. Given that free radicals (ROS and RNS) and anti-microbial peptides are primary intracellular anti-microbial factors, we analysed if these are altered in senescent cells. It has been previously observed that Reactive Oxygen Species (ROS) levels are elevated during senescence and is critical for maintenance of cell viability (Nair, Bagheri and Saini, 2014). Here also we recorded enhanced ROS levels, measured by Dichlorofluorescein diacetate (DCFDA) fluorescence and lipofuscin staining (Figs 2A and 2B). We also observed enhanced levels of another free radical species, Nitric oxide (NO), by Griess assay in S cells compared to NS cells (Fig 2C). NO production is majorly regulated by the transcript levels of *NOS2* which encodes the enzyme inducible Nitric Oxide Synthase (iNOS) that converts L-Arginine to Citrulline, and NO (Aktan, 2004). Gene expression analysis confirmed that *NOS*2 was also up regulated in S cells (Fig 2D). It is known that both nitrosative and oxidative free radicals compromise bacterial infection (Umezawa *et al*., 1997; Henard and Vázquez-Torres, 2011; Gogoi, Shreenivas and Chakravortty, 2019).

**Figure 2.**
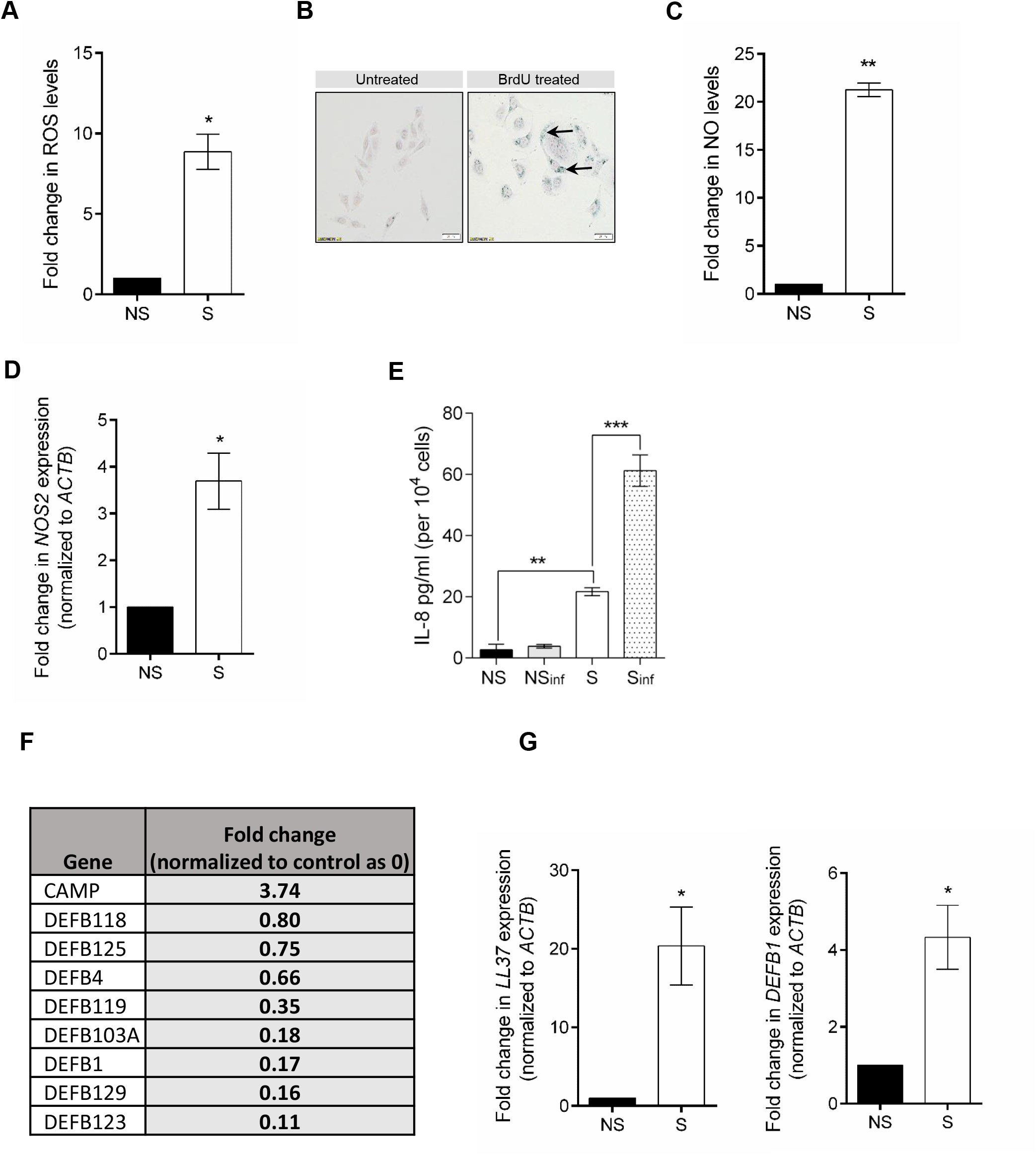
Antimicrobial defense mechanisms in senescent cells. **A**. ROS levels in NS and S HeLa cells were determined using DCFDA and normalized to levels in NS cells. **B**. Lipofuscin staining using Sudan Black B. S cale bar, 100 µm, Arrows indicate lipofuscin granules **C**. Determination of Nitric Oxide (NO) using Griess assay. **D**. *NOS2* gene expression analysis by qRT-PCR. Values were normalized to β-actin and then wrt NS cells to determine fold changes. **E**. ELISA for estimation of IL-8 secreted from uninfected and infected NS and S cells. **F**. Microarray data for expression levels changes in cationic antimicrobial peptides in senescent cells (Nair et al., 2015). Here LL37 is denoted as *CAMP*-Cathelicidin Antimicrobial Peptide and β-defensins as *DEFB* **G**. Expression analysis for LL37 and β-defensin 1 (*DEFB1*) by qRT-PCR. The data represents mean ± SEM from atleast three independent experiments. Statistical significance of differences was analysed by Mann-Whitney U test, \**P* ≤ 0.05, \*\**P* ≤ 0.01, \*\*\**P* ≤ 0.001. For all experiments NS, non-senescent and S, senescent; “inf” indicates infection.

We also estimated the levels of secreted IL-8 from NS and S cells after infection. IL-8 is a pro-inflammatory cytokine and is one of the components of the senescence associated secretory phenotype (SASP) (Coppé *et al*., 2010). Epithelial cell secreted IL-8 also acts as a chemokine for immune cell recruitment to the site of bacterial infection (Eckmann, Kagnoff and Fierer, 1993; McCormick *et al*., 1993) and hence is critical to both senescence and infection. As expected, IL-8 levels were significantly elevated in S cell secretome compared to NS cells (Fig 2E). As early as 16h after infection, IL-8 induction was observed in S cells but not NS cells (Fig 2E) indicating higher potential of senescent cells to initiate immune cell recruitment to enhance bacterial clearance.

Additionally, analysis of an unbiased microarray of NS and S HeLa transcripts reported earlier (Nair, Madiwale and Saini, 2018) suggested that the levels of cationic antimicrobial peptides (CAMPs) are also altered in senescent cells (Fig 2F). We were particularly interested in the levels of Cathelicidins (LL37) and β-defensins 1 and 2, since it has been previously reported that mice deficient for these CAMPs are susceptible to bacterial infections (Rosenberger, Gallo and Finlay, 2004; Semple and Dorin, 2012). Validation of the microarray data showed that indeed, LL37 and β-defensin 1 were also significantly up-regulated in S cells (Fig 2G), in addition to ROS and NO, however, we could not detect levels of β-defensin 2 in NS or S cells.

### Nitric oxide (NO) is the major regulator of infection in senescent cells and is negatively regulated by p38 MAPK

Compared to all other antimicrobial mechanisms, change in free radical levels in senescent cells was the most prominent and hence we decided to probe whether ROS and/or NO were important to restrict bacterial proliferation in senescent cells. For this, infections were carried out in the presence of N-acetylcysteine (NAC), a ROS quencher or Aminoguanidine (AMG), a selective inhibitor of iNOS. In the presence of the compounds, bacterial proliferation in S cells increased, however it was significantly higher in AMG treated senescent cells (Fig 3A). We further confirmed that the inhibitors did not have a direct effect at the concentrations used on *Salmonella* growth *per se* by growing the bacterium in Luria Bertani broth containing increasing concentrations of the compounds, followed by an Alamar Blue assay to assess bacterial viability. AMG did not have any direct effect at concentrations up to 1mM, however, NAC at 20µM drastically reduced bacterial viability (Fig S4) indicating a direct effect on bacterial survival. Although AMG has been extensively used as an inhibitor to iNOS, which was also confirmed by us (Fig S5A), we wanted to ensure that treatment of senescent cells with AMG does not perturb expression of other infection modulators such as anti-microbial peptides. Towards this, gene expression of the peptides, LL37 and β-defensin1 was analyzed in senescent cells after AMG treatment. Neither LL37 nor β-defensin 1 transcript levels changed (Fig S5B), demonstrating that inhibition of iNOS alone was enough to increase bacterial infection in senescent cells and CAMPs may not contribute significantly towards restricting bacterial proliferation. This highlights the importance of elevated iNOS/ NO levels in senescent cells as a major anti-bacterial factor.

**Figure 3.**
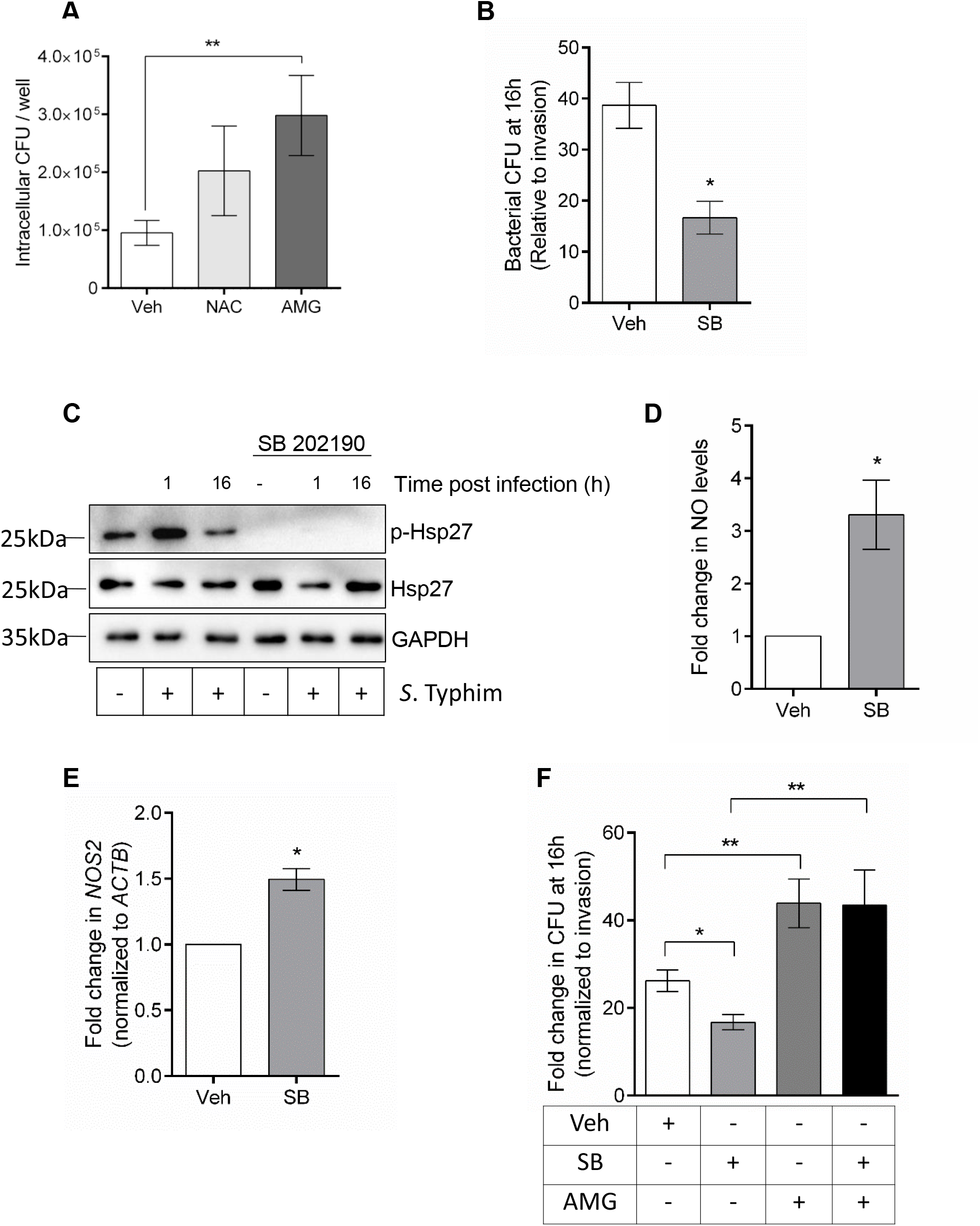
NO is the key modulator of infection and is negatively regulated by p38 MAPK in senescent cells. **A**. Bacterial CFU at 16h post invasion in vehicle, NAC and AMG treated senescent HeLa cells. **B**. p38 MAPK inhibition decreases bacterial proliferation. Senescent HeLa cells were infected in the presence of a specific p38MAPK inhibitor, SB 202190 (SB) and bacterial CFU was determined at 16 h post invasion. **C**. Western blot of p38 activity by monitoring phosphorylation status of Hsp27, a downstream substrate of p38MAPK in infected and uninfected senescent cells, treated with vehicle or SB202190. **D**. Analysis of changes in NO levels in SB202190 treated senescent cells by Griess assay. **E**. Expression analysis of *NOS2* in senescent cells treated with SB202190 by qRT-PCR. Values are normalized to β-actin and then wrt vehicle treated cells to obtain fold changes. **F**. Effect of co-inhibition of iNOS by AMG and p38 by SB202190 on intracellular bacterial proliferation. The data represents mean ± SEM from atleast three independent experiments. Statistical significance of differences was analysed by Mann-Whitney U test, \**P* ≤ 0.05, \*\**P* ≤ 0.01.

Given that NO significantly affects infection in senescent cells, we wanted to identify molecular regulators of iNOS in S cells that could affect bacterial proliferation by modulating NO levels. Since NO (Fig 2C and 2D) and inflammation (Fig 2E) are significantly higher in senescent cells, we decided to investigate the role of p38 MAPK, a well-known regulator of both inflammation and iNOS (Bhat *et al*., 2002; Cuenda and Rousseau, 2007; Freund, Patil and Campisi, 2011) in infected senescent cells. For this, senescent cells were pre-treated with SB-202190 (SB), a specific inhibitor of activated p38MAPK for 4 hours and then infection was carried out in the presence of the inhibitor. We found that bacterial proliferation was further compromised in the presence of SB-202190 (Fig 3B) without the inhibitor having a direct effect on bacterial viability (Fig S6) indicating that p38 MAPK inhibition in S cells was modulating a host anti-microbial response. Western blot for phosphorylated Hsp27 levels, a substrate of phospho-p38MAPK confirmed inhibitor activity of SB-202190 in S cells (Fig 3C). We also see that in vehicle treated senescent cells, immediately at 1h post infection, p38MAPK signaling is activated and later at 16h post infection it returns to basal levels (Fig 3C). As expected, no activation of Hsp27 was detected when infection was carried out in the presence of SB-202190.

Since the bacterial load was reduced in p38MAPK inhibited cells and from our previous findings we know that NO is a major determinant of infection, we estimated the NO levels in SB-202190 treated senescent cells using Griess assay. Indeed, NO was significantly higher in inhibitor treated cells compared to the vehicle control (Fig 3D). To further understand if transcription of iNOS was also affected, qRT-PCR was performed for its gene expression changes and we found that p38 MAPK inhibition also increased *NOS2* transcript levels (Fig 3E). When the effect of p38 MAPK inhibition on NS cells was investigated, neither infection nor NO levels were affected when NS cells were treated with SB202190 (Fig S7). Together, this indicates that p38 MAPK is a negative regulator of iNOS and therefore NO specifically in senescent cells.

To further validate that the effect on infection observed after SB 202190 treatment in senescent cells was indeed via up-regulation of iNOS, S cells were infected after co-treatment with AMG and SB 202190. As expected, treatment with SB 202190 alone decreased infection and AMG alone increased infection (Fig 3F). When both the inhibitors were used, the bacterial load was comparable to AMG treated cells (Fig 3F) and the decrease observed after SB202190 treatment was reversed. This ascertains that p38MAPK inhibition reduces infection by increasing nitrosative response of the host cell.

### Aged mice also show higher iNOS expression and reduced bacterial burden compared to young mice

It is already demonstrated that senescent cells accumulate in several tissues of old mice (Wang *et al*., 2009) and our cellular infection studies demonstrate that senescent cells significantly suppress intracellular bacterial proliferation with iNOS playing a pivotal role. Hence, to test this *in vivo*, naturally aged, male BALB/c mice (18 months old) were orally infected with *S*. Typhimurium and bacterial load in the liver and spleen, the major sites of stable bacterial colonization (Watson and Holden, 2010) was examined and compared to infected young mice (2 months old). Concurrent with our *in cellulo* findings, old mice had significantly lower bacterial load in the liver and spleen (Fig 4A). Analysis of Hematoxylin and Eosin (H&E) stained sections of the liver showed that there was recruitment of immune cells to the infected liver in both young and old mice, however, more hepatic tissue damage was seen in young infected mice (Fig 4B).

**Figure 4:**
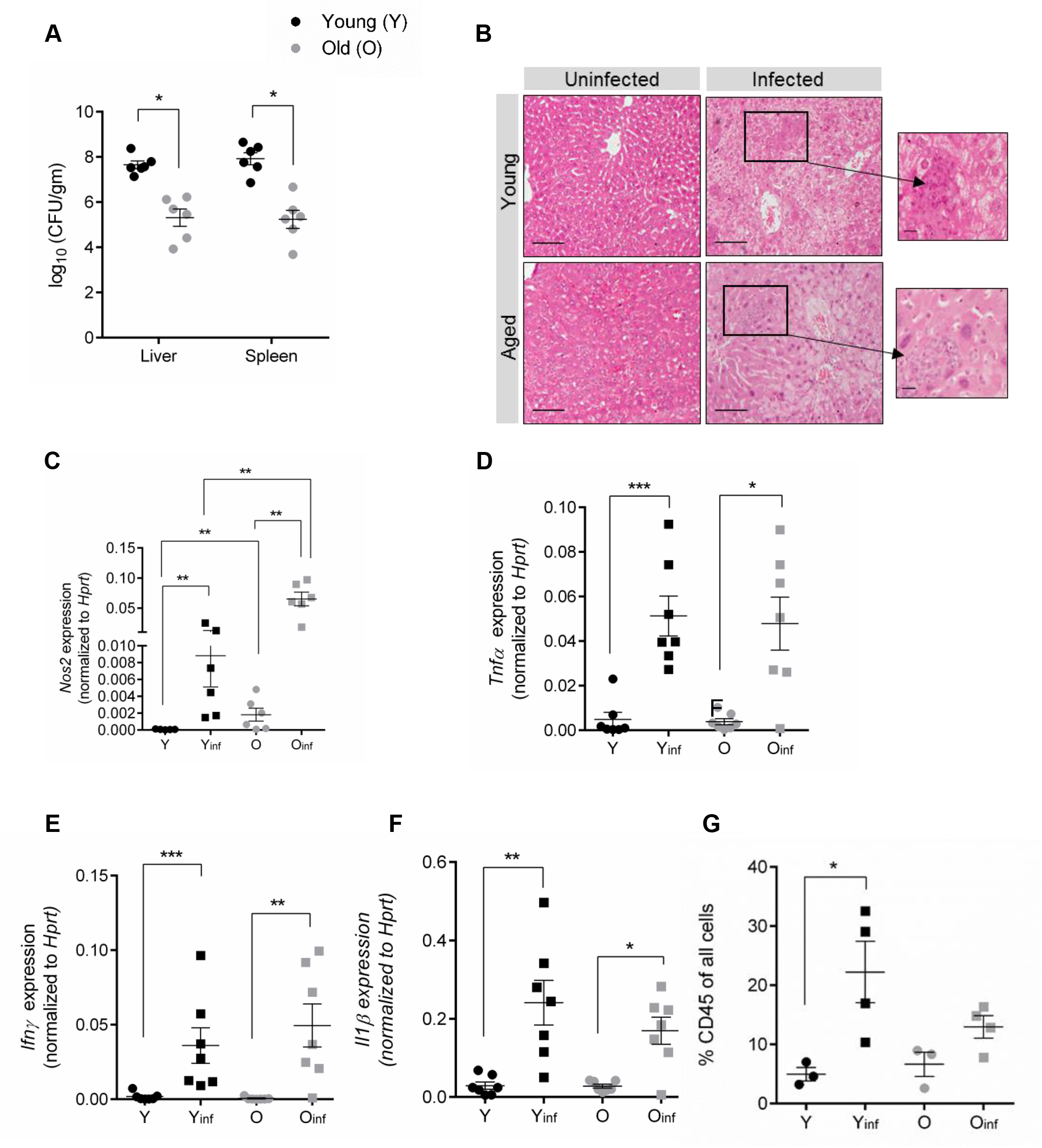
Analysis of *S*.Typhimurium infection in old and young mice. **A**. Determination of bacterial load in liver and spleen of infected mice. Young (2m old) and old (18m old) male BALB/c mice were orally gavaged with 10^8^ bacteria and CFU in the liver and spleen was determined at 4 days post infection. **B**. H&E staining of *Salmonella* infected mouse liver at 4 days post oral infection. Magnified region indicates inflammation and necrotic regions. Scale bar, 100µm. **C-F**. Gene expression analysis of *Nos2, Tnfα, Ifnγ and Il1β* from liver tissue. Values are normalized to *Hprt*. **G**. Quantification of immune cell percentages in the liver tissue of uninfected and infected mice. 4 days post infection, mice were euthanized to evaluate the immune cell presence in the liver and compared to uninfected animals. Immune cell percentages among all cells in the single suspension were ascertained by flow cytometry. (Y-young, Y_inf_-Young infected, O-Old, O_inf_-Old infected). The data shown are from individual mice, and lines depict the mean ± SEM. The differences between the experimental groups were analyzed for statistical significance using Mann-Whitney U test and one-way ANOVA followed by post-hoc Tukey test was performed for statistical analysis of immune cell data, *, *P* ≤ 0.05; **, *P* ≤ 0.01; ***, *P* ≤ 0.001.

Based on the findings from our *in-vitro* infection studies, we compared the levels of *Nos2* between young and aged mice. In agreement with our findings from senescent cells, basal *Nos2* transcript levels were higher in the liver of old mice and infection caused a further increase to levels which were significantly higher compared to young infected mice (Fig 4C). This suggests that at both cellular and organismal level of aging, increased levels of iNOS plays a significant role in restricting bacterial infection. Additionally, as expected, an increase in *Tnfα, Il1β* and *IFNγ* pro-inflammatory cytokine transcription was also observed on infection (Fig 4D-F). Since pro-inflammatory cytokines direct immune cell recruitment, we then compared the percentages of immune cells in the liver at basal level and after the mice were orally infected. We observed a significant increase in immune cells percentages in the liver of young mice after infection (Fig 4G). This corresponded with a significant increase in the percentage of monocytes and increases (but not statistically significant) in percentages of neutrophils and dendritic cells (Fig. S8B). We also observed a marginal increase in overall immune cell percentages in old mice (Fig 4G). This could be possibly due to higher bacterial burden in young mice creating the need for increased immune cell recruitment.

### Persistent inflammation compromises survival of aged mice despite lower bacterial burden

Nevertheless, when survival of mice after infection was compared by Kaplan Meier plot, old mice did not survive for a significantly longer time despite showing reduced infection (Fig 5A). This surprising observation prompted us to examine changes in the expression levels of <u>Ser</u>ine <u>P</u>rotease <u>In</u>hibitors (SERPIN) A1, B1 and B9 in the liver tissue. SERPINs are mostly known to have a protective function during chronic inflammation and an increase in their levels are correlated with resolution of inflammation induced damage to the host (Law *et al*., 2006; Choi *et al*., 2019; Kaner *et al*., 2019; Rieder *et al*., 2019). Significant downregulation of SERPINs A1 and B9 were seen in infected compared to uninfected young mice (Fig 5B and 5C) possibly to potentiate the immune response towards clearing the bacterial burden. We report a similar trend in the expression of SERPINs A1 and B9 in old mice (Fig 5B and 5C) after infection. Contrarily, SERPIN B1 levels were significantly up-regulated in young mice after infection (Fig 5D) suggesting that there is probably a balance maintained between inflammation and anti-inflammatory responses of the host to achieve clearance of pathogens while simultaneously preventing excessive damage due to the acute immune response to infection. In old mice, however, SERPIN B1 levels remained unaltered (Fig 5D) after infection indicating the lack of a protective response to infection induced inflammation. Additionally, gene expression analysis of IL-10 a known anti-inflammatory cytokine also significantly increased in young mice after infection but in old mice the increase was only marginal (Fig 5E). This suggests that the inflammatory immune response in old mice is efficient in clearing the bacteria, however, its persistently elevated levels may cause continuous damage to host tissue, compromising host survival.

**Figure 5:**
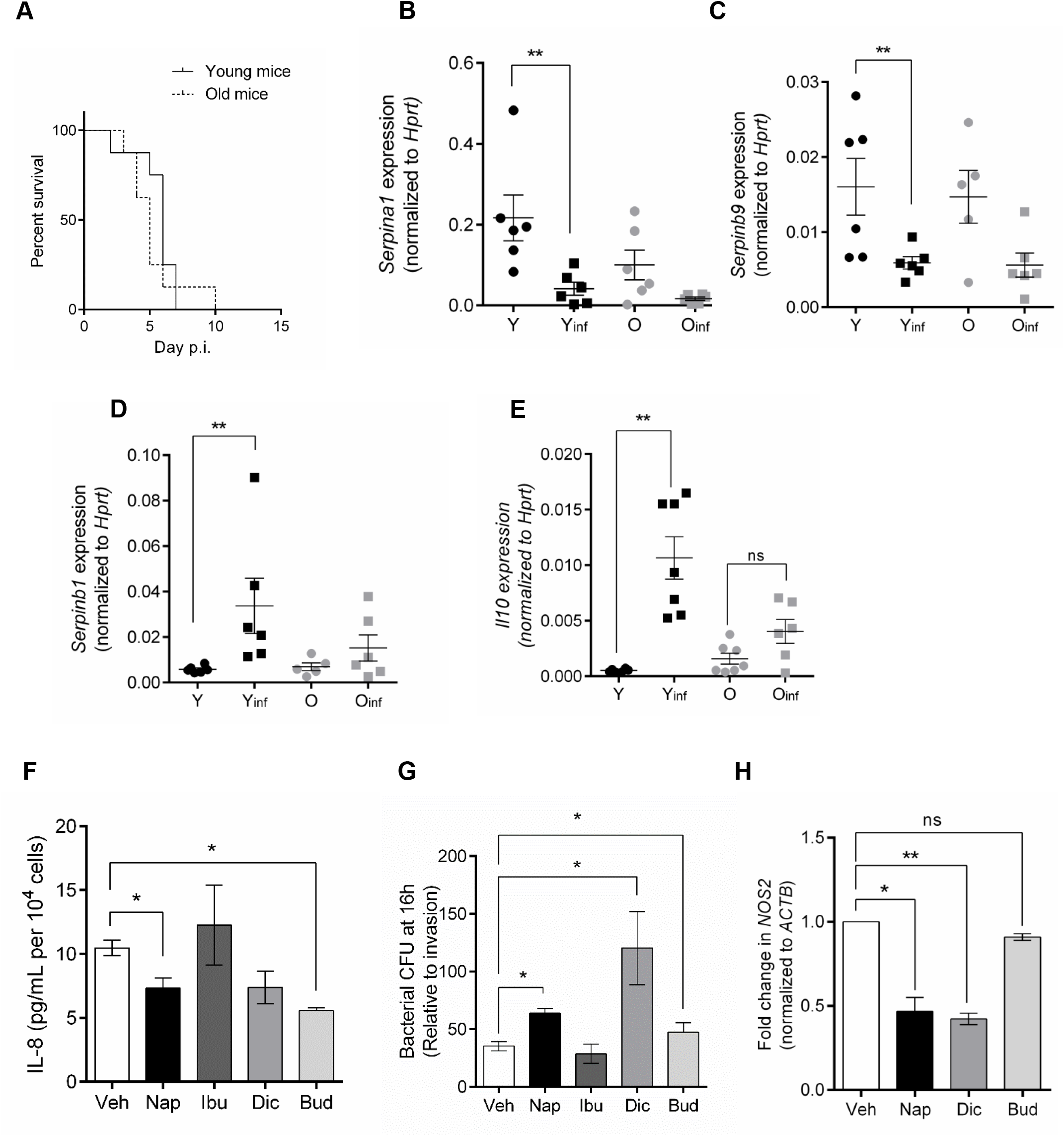
Effect of inflammation and inflammation modulating drugs on *Salmonella* infection in the aged. **A**. Kaplan Meier survival plot of young and old mice after oral infection with *Salmonella*. Mice were orally infected with 10^8^ bacteria and survival was monitored every 12 h. Each group contained at least 10 animals. **B-E**. Gene expression of various SERPINs (B-D) and IL-10 (E) in uninfected or infected mice at Day 4 post oral infection. Values are normalized to *Hprt*. The data shown are from individual mice, and lines depict the mean ± SEM. **F**. IL-8 levels measured in secretome of senescent HeLa cells treated with 10µM steroidal or NSAIDs by ELISA. **G**. Bacterial CFU at 16h post invasion in vehicle and drug treated senescent cells. **H:** Expression analysis of *NOS2* in senescent cells treated with Diclofenac, Naproxen and Budesonides by qRT-PCR. Values are normalized to β-actin and then wrt vehicle treated cells to obtain fold changes. The data represents mean ± SEM from atleast three independent experiments. Statistical significance of differences was analysed by Mann-Whitney U test, *, P ≤ 0.05; **, P ≤ 0.01; ***, P ≤ 0.001.

In fact, morbidity in the elderly is already associated with chronic inflammation and steroidal or non-steroidal anti-inflammatory drugs (NSAIDs) are commonly administered to decrease severity of several geriatric diseases (Crofford, 2013). Since our findings also reveal that inflammation can influence the survival of infected aged mice, we wanted to understand the implications of these drugs on infection in the aged. To address this, we examined effect of several common anti-inflammatory compounds viz. Diclofenac, Budesonide, Naproxen and Ibuprofen using the senescent cell model of *Salmonella* infection. We first confirmed the anti-inflammation properties of these compounds in senescent cells by analyzing the levels of secreted IL-8 after drug treatment. All the drugs except Ibuprofen were able to decrease IL-8 levels as expected (Fig 5F). However, drugs that decreased inflammation also significantly increased bacterial proliferation (Fig 5G). Having identified NO as a predominant regulator of infection, we quantified the expression of *NOS2* in the drug treated cells to find *NOS2* transcript levels significantly reduced in Diclofenac and Naproxen treated cells (Fig 5H). Increase in bacterial proliferation after Budesonide treatment was however not associated with *NOS2* suppression (Fig 5H). Thus, contrary to the expectation that NSAIDs by reducing inflammation can perhaps promote overall survival of the infected aged host, these drugs may worsen infection by compromising host anti-pathogenic responses. In support of our finding, a previous population based case-control study designed to examine the association between anti-inflammatory drugs and Non-Tuberculous Mycobacterial-Pulmonary Disorder (NTM-PD) concluded a significantly increased risk of NTM-PD in individuals being administered anti-inflammatory drugs (Brode *et al*., 2017). This study further emphasizes on the fact that administration of these drugs even as therapy for co-morbidities in the elderly may compromise important innate immune responses predisposing the elderly to infections.

### Aging associated reduction in bacterial load is also recapitulated in *Mycobacterium tuberculosis* infection

To confirm that the observed effect of aging on bacterial infection was not specific to *Salmonella*, senescent and non-senescent A549 lung epithelial cells were infected with another intracellular pathogen *Mycobacterium tuberculosis* H37Rv. Even in this model of infection, significantly lesser number of bacteria were observed at 48h post infection in senescent cells (Fig 6A). To further validate these findings at an organismal level, young and old mice were infected by aerosolization of H37Rv and bacterial burden in the lungs was determined on day 60 after infection. Similar to our observations with *Salmonella*, old mice carried significantly lesser *Mycobacteria* in their lungs compared to young mice (Fig 6B). To ensure no difference in initial bacterial loads existed between the two groups, 2 young and 2 old infected mice were sacrificed on day 1 post aerosol infection and whole lung homogenates were plated to enumerate infecting bacterial numbers. All the mice were found to be infected with≈100 bacteria and no differences were observed between the groups (Fig S8). The lungs of young mice also showed a greater number of granulomatous lesions (Fig 6C) and H&E staining of infected lung tissues revealed increased congestion and necrosis (Fig 6D) compared to old mice. The lung sections were also scored for severity of disease based on the number of granulomatous lesions, oedema and immune cell infiltration and young infected mice showed higher disease severity (Fig 6E) when compared to old infected mice. Therefore, in multiple host and intracellular bacterial pathogen models, it seems that senescence and aging reduces infection.

**Figure 6:**
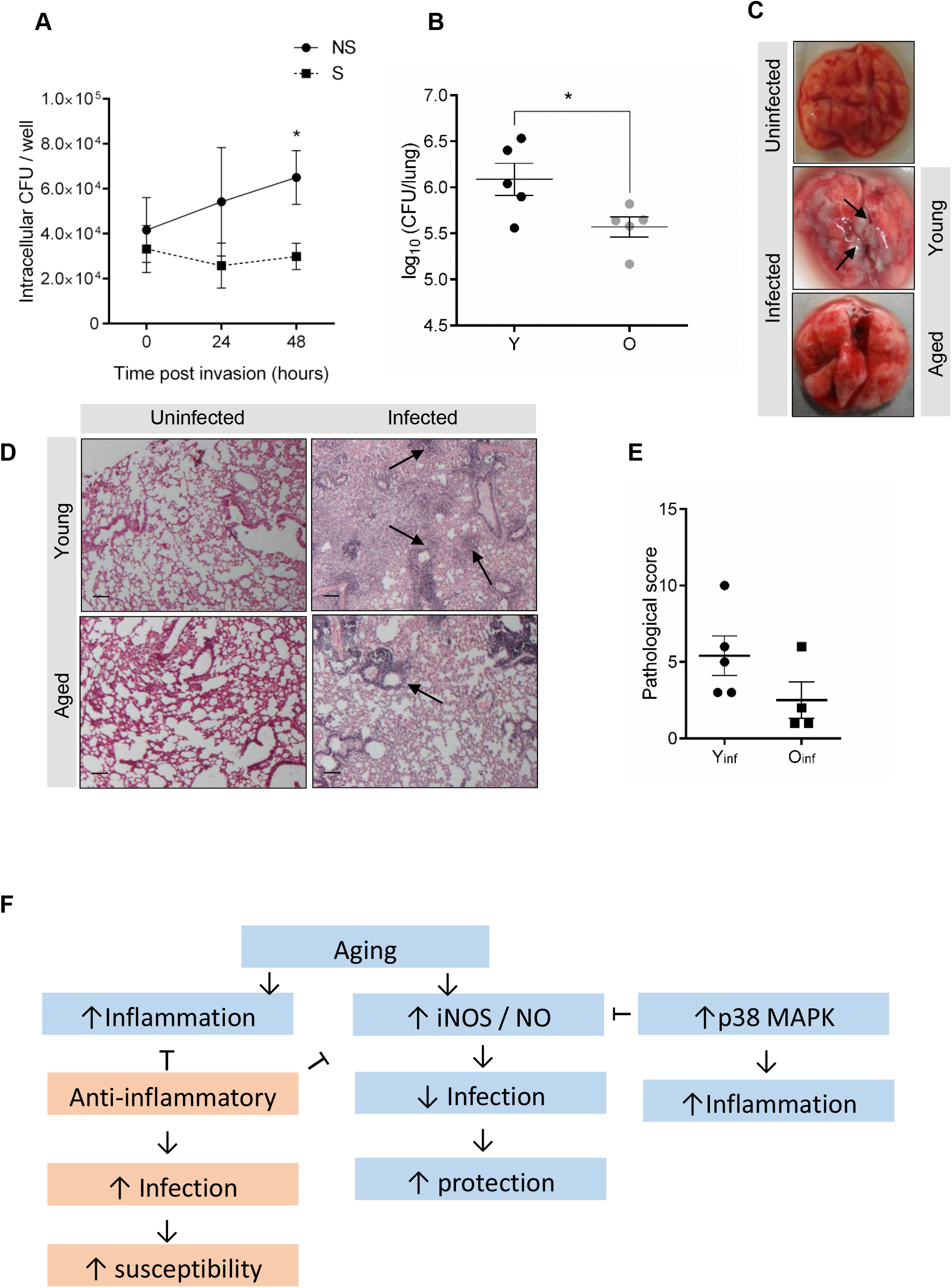
*Mycobacterium tuberculosis* infection in old and young mice. **A**. *In vitro* infection of senescent and non-senescent lung epithelial A549 cells. The data represents mean ± SEM from three independent experiments. **B**. Bacterial burden in the lungs of infected mice. Young (Y) and old (O) mice were infected *with Mycobacterium tuberculosis* (H37Rv) by aerosolization of the bacteria. At day 1 post infection, 2 mice from each group were sacrificed to ensure comparable initial bacterial load in the lungs. Subsequently, remaining infected mice were sacrificed at day 60 post infection to determine bacterial load in the lungs. The data show the values for individual mice, with error bars showing the mean ± SEM. **C**. Lungs from infected mice showing oedema and granulomatous lesions (Arrow). **D**. H&E staining of *Mycobacterium* infected lung tissues. Arrows indicate necrotic, granulomatous lesions and congestion. **E**. Pathological scoring of lung tissue sections of infected mice based on number of granulomatous lesions, oedema and immune cell infiltration. Scale bars, 100µm. The statistical significance of the differences among experimental groups in all panels was analysed using Mann-Whitney U test, \**P* ≤ 0.05. **F**. A model with the salient findings of the study.

## Discussion

Accumulation of senescent cells in various tissues of the body is the major driver of aging (Jeyapalan *et al*., 2007) and is associated with several geriatric metabolic and degenerative disorders (Baker *et al*., 2011; Campisi and Deursen, 2016; Jeon *et al*., 2017) but there is a paucity of literature that focusses on how it may affect infections in the elderly. In a study by Shivshankar *et al*, it has been reported that increase in the expression of the receptor K10, specific to keratinocytes and mucosal epithelial cells of the lungs increases adhesion of the extracellular pathogen *S*.*pneumoniae*, and could possibly be responsible for higher incidence of pneumonia seen in the elderly (Shivshankar *et al*., 2011). Another study implicates age associated Monocyte dysfunction for the reduced anti-pneumococcal activity observed in the aged (Puchta *et al*., 2016). Many other studies on bacterial infection have focussed on how changes in the immune system during ageing can influence infection. However, contradicting evidence regarding immune-senescence indicates the presence of other confounding factors (Esposito and Pennington, 1983; Cooper *et al*., 1995; Pacheco *et al*., 2013), which may decide the final outcome of infection in the aged.

In the present study, we have used both cellular and organismal models to elucidate the effect of aging on two intracellular bacteria *S*.Typhimurium and *Mycobacterium tuberculosis*. Lim *et al* have previously reported that invasion of *S*.Typhimurium is enhanced in senescent fibroblast cells compared to non-senescent fibroblasts, however they do not comment on bacterial proliferation or survival post invasion (Lim *et al*., 2010). In contrast to their study, using established epithelial cell models as hosts, specifically HeLa for *Salmonella* Typhimurium and A549 for *Mycobacterium tuberculosis*, we show that the ability of bacteria to invade non-senescent or senescent epithelial cells does not vary in contrast to fibroblasts, but senescent cells significantly inhibit intracellular bacterial proliferation. Investigation of known antimicrobial factors that could contribute to this phenotype reveal that several antibacterial mechanisms like levels of free radicals and antimicrobial peptides are already elevated in senescent cells possibly allowing them to respond to invading bacteria more rapidly than non-senescent cells. However, of all the enhanced mechanisms, elevated levels of Nitric Oxide (NO) was found to play a pivotal role in limiting bacterial proliferation. Mechanistically, we demonstrate that p38 MAPK in senescent cells keeps NO levels in check and its inhibition causes increase in the transcription of *NOS2* which encodes the enzyme inducible <u>N</u>itric <u>O</u>xide <u>S</u>ynthase (iNOS) responsible for the production NO. Inhibition of p38 MAPK was able to increase NO to levels that could further reduce bacterial infection indicating a key role of p38 MAPK in regulating Nitric Oxide levels in senescent cells and thereby infection.

The relevance of our findings from cellular infection studies were also tested *in vivo* using naturally aged mice. Previously using the Streptomycin induced gastroenteritis mouse model, it was demonstrated that old mice had higher tissue colonization and morbidity compared to young mice (Ren *et al*., 2009). However, Streptomycin induced colitis may also directly affect infection outcomes. From our infection studies, we see that old mice had significantly reduced bacterial burden in the lungs upon aerosol infection with *M*.*tuberculosis* or in the liver and spleen when orally infected with *S*.Typhimurium. Importantly, iNOS transcript levels were higher in liver tissue of old animals and was further increased upon infection, thus re-enforcing our conclusions from *in vitro* infection experiments that elevated levels of NO during aging may play an important role in modulating infection.

Despite the lower bacterial infection, survival of old mice did not significantly vary from young mice, possibly because mortality is a cumulative outcome of the host’s ability to fight the pathogen while simultaneously protecting itself from inflammation induced tissue damage. Here we show that infected old mice mount significant anti-pathogen responses including increase in pro-inflammatory cytokines TNFα, IL1β and IFNγ and NO generating NOS2 enzyme, but made no attempt to up-regulate anti-inflammatory mediators such as IL-10 and SERPINs resulting in lower bacterial loads but persistent inflammation. In fact, elevated inflammation in the elderly has been previously associated with enhanced mortality (Giovannini *et al*., 2011; de Gonzalo-Calvo *et al*., 2012) and reduced survival after pneumococcal infection (Yende *et al*., 2013). Contrastingly, young mice elicited a concomitant anti-inflammatory response but these together with a reduced anti-bacterial nitrosative response favored bacterial survival and proliferation. Overall, this suggests that mortality of young mice maybe a result of bacterial infection but that of old mice maybe a result of its own inflammatory response and not due to the bacterial loads *per se*.

Apart from affecting survival after infection, chronic inflammation also increases the severity of age associated co-morbidities. Hence, several anti-inflammatory compounds are being administered (Walker and Lue, 2007; van Walsem *et al*., 2015) and other compounds re-purposed under the broad category of Senotherapeutics to reverse the deleterious effects of inflammation. However, we show that common FDA approved anti-inflammatory drugs like Naproxen, Diclofenac and Budesonide which maybe recommended also enhanced bacterial survival. Therefore, from an infection perspective, although these drugs may help to decrease levels of pro-inflammatory factors, they are likely to parallelly create conducive conditions for bacterial survival and proliferation by interfering with innate cellular immune responses like NO production.

In summary (Fig 6F), we demonstrate that several innate anti-microbial factors are elevated in senescent epithelial cells and old mice. The result of these changes is an overall reduced bacterial burden upon infection both *in vitro* and *in vivo* and highlights an alternate advantage of senescent cell accumulation in the elderly who are witnessing a simultaneous decline in adaptive immunity. Further, therapies recommended to treat co-morbidities in the elderly may compromise these factors, thereby increasing susceptibility of the elderly to infection.

## Materials and Methods

### Cell culture and induction of senescence

HeLa, HepG2 and A549 (ATCC, USA) were maintained in DMEM (Sigma Aldrich, USA) supplemented with 10% FBS (Invitrogen). Senescence was induced by treating cells with 100 µM 5-Bromo-2’-deoxyuridine (Sigma Aldrich, USA) for 48 hours. HepG2 cells were a kind gift from Prof. Saumitra Das, Department of Microbiology and Cell Biology, IISc, Bangalore.

### Bacterial infections in cells

For Salmonella infections, the *S*. Typhimurium NCTC 12023 strain was used. An overnight culture prepared from a single isolated colony of *S*. Typhimurium grown on a *Salmonella-Shigella* (SS) agar plate was diluted 1:200 in LB and grown for 6 h at 37°C and 180 rpm to obtain a log-phase culture (OD_600_ 1.0). The bacterial culture was then washed and resuspended in sterile phosphate-buffered saline (PBS) and used for infection.

To quantify bacterial infection, a monolayer of non-senescent or senescent HeLa/HepG2 cells was infected at MOI 1:10 for 60 minutes (or for 30 minutes to quantify invasion differences) at 37°C in a 5% CO2 humidified atmosphere (invasion). Immediately after addition of bacteria, cells were centrifuged at 250×g for 10 minutes at room temperature (RT) to allow synchronous invasion of cells. At the end of co-incubation, the medium was replaced with fresh complete medium containing 100 µg/mL Gentamicin for 30 minutes to kill extracellular bacteria. Cells were then lysed in 0.5% Triton X-100 (v/v in PBS) to enumerate the number of bacteria that have invaded or maintained in medium supplemented with 10 μg/ml gentamicin until further time points at which they were lysed to determine intracellular bacterial survival and proliferation after invasion. Dilutions of the lysates were plated on SS agar to enumerate bacterial CFU.

For imaging, infections were carried out as mentioned above but using mCherry expressing S. typhimurium 12023. At 16 hours post invasion, the cells were fixed with 4% paraformaldehyde and imaged using an Olympus IX83 inverted fluorescence microscope.

To quantify the percentages of cytosolic *Salmonella*, a Chloroquine (CHQ) resistance assay was performed as previously described by Knodler *et al* (Knodler, Nair and Steele-mortimer, 2014). Briefly, cells were infected as mentioned above and 1h prior to lysis, CHQ at a concentration of 400µM was added to the gentamicin containing medium, to determine CHQ resistant and hence cytosolic bacteria. Two other wells were simultaneously maintained without CHQ (total bacteria). Cells were then washed, lysed and dilutions of the lysates were plated on SS agar. Percent cytosolic bacteria was calculated as (bacterial CFU after CHQ treatment/bacterial CFU without CHQ treatment) *100.

To study infection in the presence of molecular inhibitors or drugs, the cells were pre-treated with compound for 4 hours prior to infection and the compounds were maintained in the medium during infection till cells were used for further analysis. Aminoguanidine hemisulfate salt (AMG) (Sigma Aldrich, USA), N-acetyl-L-Cysteine (Sigma Aldrich, USA), p38 MAPK inhibitor SB 202190 (Cayman Chemical Co., USA), Diclofenac (Cayman Chemical Co., USA), Budesonide (Cayman Chemical Co., USA), Naproxen (Cayman Chemical Co., USA) and Ibuprofen (Cayman Chemical Co., USA) were used at a concentration of 10µM unless specified otherwise.

For *Mycobacterium tuberculosis* infections, an actively growing culture of virulent strain *Mycobacterium tuberculosis* H37Rv in 7H9 broth supplemented with 10% OADC was used to infect a monolayer of non-senescent or senescent A549 cells at MOI 1:10 for 4 h at 37°C in a 5% CO_2_ humidified atmosphere. The cells were then washed with PBS, to remove extracellular bacteria and lysed in 0.5% Triton X-100 (v/v in PBS) or maintained in complete medium until further analysis. CFU was determined by plating the lysate on 7H11 plates supplemented with 10% OADC.

### Animal infections

Young (2 months) and old (18 months) male, BALB/c mice were obtained from Government establishment. For *Salmonella* infections, the mice were orally gavaged with 10^8^ bacteria resuspend in 200µL of PBS. At 4 days post infection, the mice were sacrificed, and the liver and spleen were harvested for CFU determination, gene expression and histological analysis.

For *Mycobacterium tuberculosis* infections, the mice were aerosolized with 500 CFUs of Mtb H37Rv and maintained in securely commissioned BSL3 facility for 60 days. The animals were then euthanized, and lungs were harvested for CFU determination and histological analysis.

### Ethics statement

The experiments were performed in agreement with the Control and Supervision Rules, 1998 of Ministry of Environment and Forest Act, and the Institutional Animal Ethics Committee and experimental protocols were approved by the Committee for Purpose and Control and Supervision of Experiments on Animals (permit number CAF/Ethics/588/2018).

### Lipofuscin staining

Lipofuscin staining was essentially done as described by Georgakopoulou *et al* (Ea *et al*., 2013). Cells seeded on coverslips were fixed in 4% PFA and followed by incubation in 70% ethanol for 2 min. Coverslips were then inverted on a slide containing a drop of Sudan Black B solution (0.7 gram dissolved in 100mL 70% ethanol). Excess stain was washed away using 50% ethanol followed by distilled water and cells were then counterstained with 0.1% Eosin.

### Gene expression analysis

Total cellular RNA was isolated using TRI reagent (Sigma, USA) and cDNA was synthesised using iScript cDNA Synthesis Kit (Bio-Rad, USA) followed by quantitative expression analysis using SYBR Green qPCR Kit (Thermo Fisher Scientific, USA) as per manufacturer’s instructions. Expression levels of β-actin and HPRT were used to normalize the expression levels in cells and animal tissues respectively. RotoGene-Q real-time instrument and associated software was used for data and melting curves analysis. Primers used are mentioned in Table S1.

### SA-β gal staining

The protocol described by Dimri et al. was followed for SA β-gal staining (Dimri et al. 1995). Cells were washed, fixed for 15 minutes at room temperature in 0.2% glutaraldehyde (Amresco, USA) prepared in PBS and then incubated overnight at 37°C (without carbon dioxide) with freshly prepared staining solution (1mg/ml of X-gal (GoldBio Technology, USA) in 40 mM citric acid/sodium phosphate, pH 6.0, 5mM potassium ferrocyanide, 5 mM potassium ferricyanide, 150 mM NaCl and 2 mM magnesium chloride. Cells were then washed with PBS to get rid of excess stain and imaged using an inverted IX81 microscope, equipped with DP72 colour CCD camera (Olympus, Japan).

### Western blotting

Cell lysates were prepared in ProteoJET Mammalian Cell Lysis Reagent (Fermentas Inc., USA) as per manufacturer’s specifications. 60-100µg of total protein was used for analysis. All the primary antibodies were from CST (Cell Signalling Technology Inc., USA) and used at 1:1000 dilution overnight at 4°C for probing protein levels viz. phospho-Hsp27 (Cat No. 9709), Hsp27 (Cat No. 95357) GAPDH (Cat No. 2118) and phospho-Chk2 (Cat no. 2197). The developed blots were imaged and analysed using ChemiDoc MP Imaging system (Bio-Rad Inc., USA) at multiple exposure settings.

### ROS estimation

Cells were incubated with 10 µM 2′,7′-dichlorofluorescein (DCFDA) (Sigma, USA) in PBS for 30 min in dark, washed and analysed to detect DCF fluorescence (Infinite F200, Tecan, Austria) at an excitation wavelength of 492 nm and emission wavelength of 525 nm. Cells were counted to express DCFDA fluorescence per cell.

### NO quantitation

Griess reagent (Sigma, USA) was used to measure nitrite as an indicator of NO, according to manufacturer’s protocol. Briefly equal volumes of cell supernatant (50 μl) and Griess reagent were mixed in a 96-well flat-bottom microtiter plate and absorbances were read at 550 nm using a microtiter plate reader (Tecan, Austria). The amount of NO produced was determined using a standard curve for nitrite (1.56-100 μM NaNO_2_).

### Immune cell profiling

Single-cell suspensions from the liver were prepared in PBS containing 1 % BSA and 4 mM EDTA. Cells were stained with a combination of the following antibodies for 30 min at 4°C in presence of Fc Block: Ly6G (clone 1A8), Ly6C (AL-21), CD11b (M1/70), CD11c (HL3), F4/80 (T45-2342), CD45 (30-F11) all purchased from BD Biosciences (San Diego, CA, USA). Live cells were identified by staining cells with propidium iodide (2µg /ml) and applying a negative gating strategy. Appropriate fluorescence-minus-one (FMO) controls were used to gate positive populations. Flow cytometry data were collected using a BD FACS Celesta and analyzed using FlowJo (Tree Star, Ashland, OR, USA).

### ELISA for IL-8

Extracellular levels of IL-8 were estimated using BD OptiEIA™ Human IL8 ELISA kit (BD Biosciences, USA) as per manufacturer’s instructions. Media was collected from treated cells as indicated and cells were counted to normalize the IL-8 concentrations determined to 10^4^ cells. It was ensured that the raw values obtained were within the dynamic range of the assay.

### Statistical analysis

For cell-based experiments, biological triplicates or more were used. And for animal experiments 5 or more animals were used per group. All n’s are mentioned in the figure legends. Prism software (GraphPad Prism 6.0) was used for the generation of graphs and analysis. For all experiments, results are represented as mean ± SEM. For statistical analysis, the Mann–Whitney test was used for the comparison of medians from two groups or One-way ANOVA followed by post-hoc Tukey test for comparison of multiple groups. Significance (p value) is represented as *, where *≤0.05, **≤0.01, ***≤0.001, and ****≤0.001 and ns, where >0.05 for “not significant”.

## Acknowledgments

We are grateful to Prof. Dipankar Nandi for his suggestions and inputs and Central Animal Facility (CAF), IISc for the animal work. We also acknowledge Dr. V. Ravikumar (RV Metropolis, Bangalore) who assisted in histopathological analysis.

## Data availability

The data that support the findings of this study are available from the corresponding author upon reasonable request.

